# Environmental deformations dynamically shift the grid cell spatial metric

**DOI:** 10.1101/174367

**Authors:** Alexandra T Keinath, Russell A Epstein, Vijay Balasubramanian

## Abstract

Environmental deformations induce stereotyped distortions in the time-averaged activity of grid and place cells. We hypothesized that these effects are partly driven by border cell inputs which reset the spatial phase of grid cells, maintaining learned relationships between grid phase and environmental boundaries without altering inherent grid scale. A computational model of this mechanism reproduced diverse distortions during deformations, including scale-dependent and local distortions of grid fields, and stretched, duplicated, and fractured place fields. This model predicted a striking new effect: dynamic, history-dependent, boundary-tethered ‘shifts’ in grid phase during deformations. We reanalyzed two rodent grid cell rescaling datasets and found direct evidence of these shifts, which have not been previously reported and contribute to the appearance of rescaling. These results demonstrate that the grid representation of geometrically deformed environments is not fixed, but rather dynamically changes with the specific experience of the navigator.

---

The cognitive map is thought to be a metric representation of space that preserves distances between represented locations [1,2]. Entorhinal grid cells are hypothesized to generate this metric by maintaining an internally-generated, path-integrated representation of space [3–8]. Results of environmental deformation experiments have led to the belief that this metric is fundamentally malleable [9–12]. In these experiments, neural activity is recorded as a rat explores a familiar environment that has been modified by stretching, compressing, or removing/inserting chamber walls. Such deformations induce a number of distortions in the time-averaged activity of both grid cells [9,11] and hippocampal place cells [13–17]. Often described as ‘rescaling’, these distortions have been taken to suggest that the spatial metric of the cognitive map can be reshaped by altering environmental geometry [9,18,19]. Crucially, this interpretation assumes that the distortions observed in the time-averaged rate maps of these cells reflect fixed changes to the underlying spatial code that are independent of the movement history of the navigator. Here, we present results that challenge this assumption, and indicate the grid cell spatial metric undergoes dynamic history-dependent phase shifts during environmental deformations.

Our treatment focuses on the contribution of border cell-grid cell interactions to deformation-induced grid and place cell distortions. Border cells, co-localized with grid cells in the entorhinal cortex, are active only when a boundary is nearby and at a particular allocentric direction [20,21], similarly to boundary vector cells [22]. Stretching or compressing a boundary yields a concomitant rescaling of border activity neighboring that boundary, and insertion of a new boundary elicits additional border activity at analogous locations neighboring the new and old boundaries. In familiar undeformed environments, input from border cells is thought to a correct drift in the grid pattern [23,24], and it has been suggested that input from border cells may influence the activity of grid and place cells during environmental deformations [10,20,23,25–27]. However, the ways in which border cell-grid cell interactions might shape grid and place cell activity during deformations have not been fully characterized and specific experimental evidence of such a contribution is lacking.

To address this question, we first constructed a model where the activity of a grid cell attractor network [28] is shaped by Hebbian-modified input from border cells [20]. The model also included a population of units corresponding to hippocampal place cells, whose responses were learned from grid unit output [29,30]. Our simulations showed that during environmental deformations, this model reproduces a number of experimentally-observed phenomena: (1) when a familiar environment is rescaled, the firing patterns of large-scale grid units rescale to match the deformation, while the firing patterns of small-scale grid units do not [9,11]; (2) when a familiar environment is partially deformed, the neighboring grid structure is locally distorted [12]; (3) when a familiar environment is stretched, the fields of place units exhibit a mix of stretching, bifurcation, modulation by movement direction, and inhibition [13]; (4) when a familiar linear track is compressed, the place code is updated when a track end is encountered [14,31]; (5) when a new boundary is inserted in an open environment, place fields exhibit a mix of duplication, inhibition, and perseverance [15–17]. This model further generated a striking new prediction: grid fields should exhibit shifts in grid phase that are dependent on the most recently contacted boundary, an effect we term *boundary-tethered shift*. To test this prediction, we reanalyzed datasets from two previous environmental deformation experiments [9,11], and found previously unnoticed evidence of boundary-tethered phase shifts in recorded grid cell activity. Together, these results indicate that geometric deformations of a familiar environment induce history-dependent shifts in grid phase, and implicate border cell-grid cell interactions as a key contributor to deformation-induced grid and place cell distortions.

## Results

### A model of border, grid, and place cell interactions

We implemented a spiking model of the interactions between border, grid, and place cells as follows. The border population consisted of 32 units whose activity was designed to mimic the behavior of border cells [20]. (Throughout this paper, we use ‘unit’ to refer to modeled data, and ‘cell’ to refer to *in vivo* recorded data.) Each border unit was active only when a boundary was nearby, within 12 cm in a particular allocentric direction [23]. The preferred firing field of each border unit covered 50% of the perimeter length, and maintained proportional coverage if that boundary was deformed [20,21,24] (Fig. 1). Border fields were uniformly distributed around the perimeter of the environment. If a new boundary was inserted, the border unit was active at an allocentrically analogous location adjacent to the new boundary [20,21].

**Figure 1.**
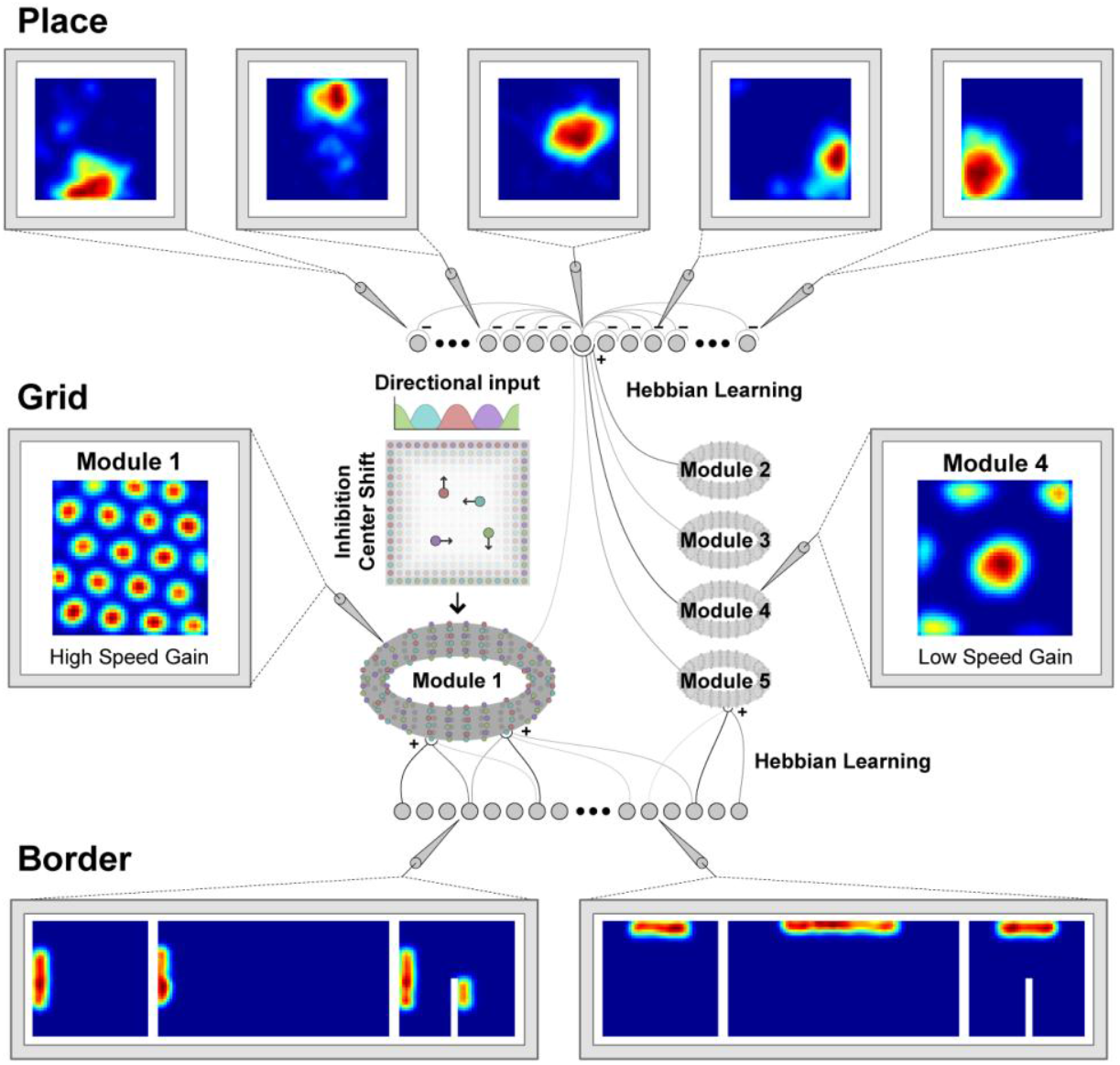
*Schematic of the boundary-tethered model network*. *The network model consisted of three layers: a border layer, where unit activity was determined by the presence of a boundary nearby and in a particular allocentric direction; a grid layer, where path integration implemented by a periodic attractor network of the form described in [28] was used to generate 5 modules of grid units of different scales; and a place layer, where unit activity was learned from the output of grid units of all scales in concert with recurrent inhibition. Excitatory connections from border cells to grid cells were learned with experience in the familiar environment. Border fields are taken to stretch when their preferred boundary is stretched and duplicate with a similar allocentric relationship to both boundaries when a boundary is inserted*.

The grid population was subdivided into 5 modules, each consisting of a neural sheet of size 128 × 128 units. The internal connectivity and dynamics of each module was based on the attractor network model described in [28], and was identical across modules except for a single movement velocity gain parameter controlling the grid scale of each module. This parameter was adjusted to yield a geometric series of scales across modules (scale factor of 1.42), as observed experimentally [11] and explained theoretically [32,33]. In addition to these connections, each grid unit also received initially random excitatory input from all border units. These connections developed through experience via a Hebbian learning rule in which connections between coactive grid and border units were strengthened at the expense of connections from inactive border units [34].

The place population consisted of 64 units receiving initially random excitatory input from 500 random grid units. These connections also developed with experience via Hebbian learning [30,34]. In combination with uniform recurrent inhibition, these dynamics yield place-cell-like activity at the single unit level.

### Model grid units deform with the environment in a scale-dependent and local fashion

Electrophysiological experiments have shown that rescaling a familiar environment can induce a corresponding rescaling of grid cell firing patterns, dependent on grid scale [9,11]. To explore the effects of environmental rescaling on grid units, we first familiarized a naive virtual rat with a 150 cm x 150 cm square environment. During this familiarization period, the border-grid connectivity self-organized via Hebbian learning (see Materials and Methods). The virtual rat then explored the familiar environment and deformed versions of this environment without new learning (chamber lengths between 75 cm to 225 cm in increments of 25 cm; chamber sizes chosen to match experiment [11]). Consistent with previous reports [9,11], we observed that these deformations induced rescaling of time-averaged rate maps in some grid modules (Fig. 2A). To quantify this module-dependent rescaling, we computed the grid rescaling factor required to stretch or compress the time-averaged rate maps in the familiar environment to best match the rate maps in the deformed environment, separately for each module. We found that the grid patterns of units in large-scale modules morphed with the environment, but grid patterns of units in small-scale modules tended not to (Fig. 2B). Precisely this behavior is observed experimentally [11]. These results demonstrate that input from border cells is sufficient to induce scale-dependent grid rescaling.

**Figure 2.**
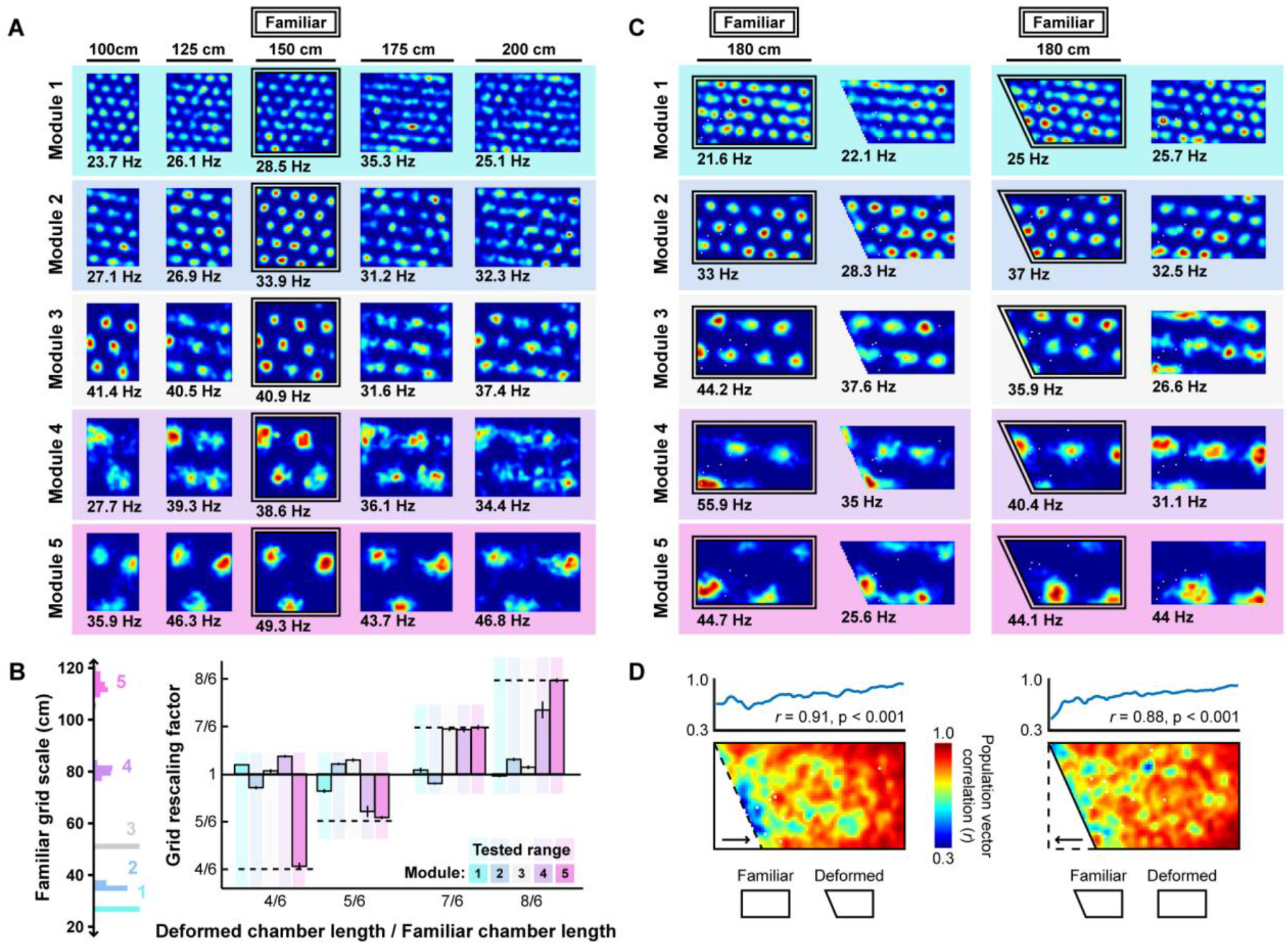
*Grid unit responses to deformations of an open environment*. *A) Rate maps from one grid unit from each module across all rescaling deformations. Colors normalized to the maximum across each set of rate maps. Peak firing rate for each trial noted below the lower left corner of each map. B) Grid rescaling factors for each module when the familiar open environment is rescaled to various chamber lengths (right). Error bars denote standard error of the mean (SEM) across 30 random grid units. Color denotes module. Distribution of grid scales for each module indicated (left). C) Rate maps of one grid unit from each module across each partial deformation, plotted as in (A). D) Correlation between the familiar and deformed environment rate maps across the population (150 grid cells, 30 random cells from each module) at each location (bottom) and averaged across north-south positions (top)*.

Next, we explored how partial deformations affect model grid units. Recording experiments have demonstrated that displacement of part of one wall of a familiar environment distorts the grid pattern locally near that wall, with neighboring grid fields shifting in the displaced direction [12]. We first familiarized a naive virtual rat with either a 180 cm x 90 cm rectangular or right trapezoid environment (long parallel wall of the right trapezoidal environment was 180 cm, short parallel wall was 135 cm, Fig. 2C). During this familiarization period, the border-grid connectivity self-organized via Hebbian learning. Without new learning, the rat then explored both the rectangular and right trapezoid environments. During deformations, fields near the displaced wall were distorted, often shifting in concert with the displaced wall, while fields far from this wall were less affected (Fig. 2C). To quantify this pattern, we computed the correlation between the familiar and deformed environment rate maps across the population at each location, sometimes called the population vector correlation. This correlation was high at locations far from the displaced wall, but was reduced near the displaced wall (Fig. 2D). Thus, border cell-grid cell interactions can give rise to local distortions similar to those observed experimentally during partial deformations. Together, these results demonstrate that many of the complex grid distortions observed during environmental deformations can emerge from border cell-grid cell interactions.

### Model place units deform heterogeneously during environmental deformations

Electrophysiological experiments have shown that stretching a familiar environment induces a heterogeneous mix of responses in the time-averaged activity of place cells [13]. To explore the effects of stretching deformations on model place units, we began by familiarizing the naive virtual rat with a 61 cm x 61 cm square open environment, during which period the border-grid connectivity and grid-place connectivity self-organized via Hebbian learning. Following this familiarization, the virtual rat again explored the familiar environment, as well as a number of deformed environments without new learning (various chamber lengths between 61 cm and 122 cm, chamber widths 61 cm or 122 cm; chamber sizes chosen to match experiment [13]). During these deformations, we observed heterogeneous changes in the time-averaged rate maps of place units (Fig. 3A). A number of place units exhibited place field stretching in proportion to the rescaling deformation. Other units exhibited place field bifurcations accompanied by progressively lower peak firing rates during more extreme deformations. Finally, some units exhibited emergent modulation by movement direction, with place fields shifting ‘upstream’ of the movement direction. A qualitatively similar mix of place field distortions is observed experimentally [13].

**Figure 3.**
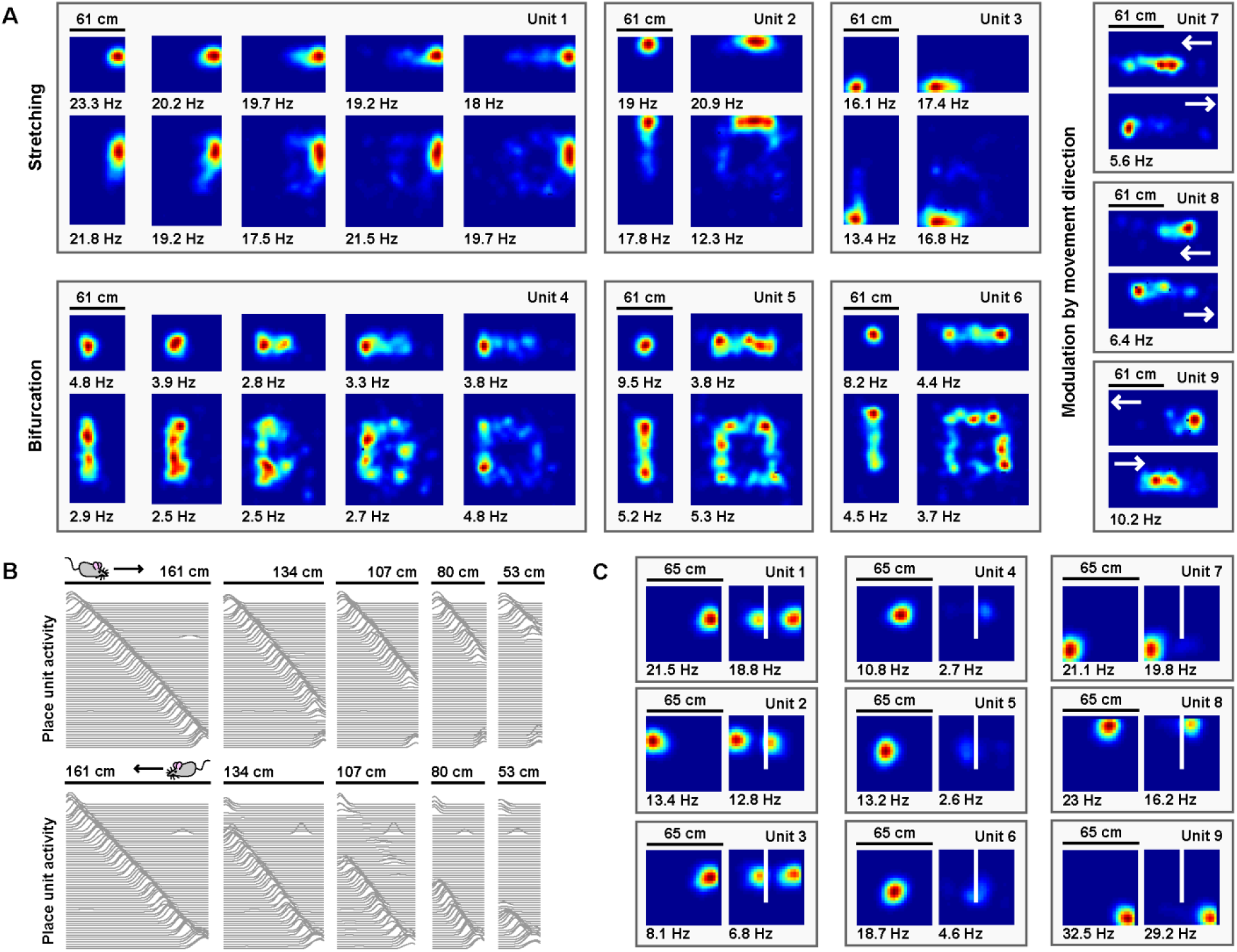
Place unit responses to deformations of open and linear track environments. *A) Place unit rate maps when a familiar open environment is stretched. Place fields exhibit stretching, bifurcation, and emergent modulation by movement direction (indicated by white arrows). Colors normalized to the peak for each rate map. Peak firing rate noted below the lower left corner of each map. Note that peak firing rate tends to decrease with more extreme deformations for cells with place fields further from boundaries. B) Place unit activity for all 64 place units during compressions of a familiar linear track, separated by (top) eastward and (bottom) westward laps. Each line indicates the firing rate of a single place unit at each location across the entire track during movement in the specified direction, normalized to the familiar track peak rate. Units sorted by place field location on the familiar track. Note that, during compressions, the place code unfolds as if anchored to the beginning of the track until the end of the track is encountered, at which point the familiar end-of-track place units are reactivated. C) Place unit rate maps demonstrating a mix of place field (left) duplication, (middle) inhibition, and (right) perseverance when a new boundary (white line) is inserted in a familiar open environment. Colors normalized to the maximum of both rate maps. Peak firing rate noted below the lower left corner of each map*.

Electrophysiological experiments have also demonstrated that when a familiar linear track is compressed, the place code is updated when track ends are encountered [14,31]. We therefore examined the effects of compressing a familiar linear track on model place units. We first familiarized the naive virtual rat with running laps on a 161 cm long linear track, during which period the border-grid connectivity and grid-place connectivity self-organized via Hebbian learning. Following this familiarization, the virtual rat ran laps along both the familiar track and a number of compressed tracks without new learning (track lengths between 53 cm to 161 cm; lengths chosen to match experiment [14]). During laps on compressed tracks, place unit activity unfolded as if unaffected by the compression, no matter how extreme, until the opposing track end was reached. Once encountered, the place code previously active at this track end during familiarization reemerged (Fig. 3B), as observed experimentally [14]. In recording experiments, similar boundary-tethered updating persists in darkness indicating that such dynamics arise even in the absence of visual cues [31], a result consistent with the sustained activity of border cells in darkness [35,36]. However, we note that in these recording experiments the particular transition point differs depending on the availability of visual input and may precede border cell firing, which likely reflects the influence of additional mechanisms outside the scope of our boundary-tethered model [18,19].

Finally, electrophysiological experiments have shown that when a boundary is inserted in a familiar open environment, place fields exhibit a mix of duplication, suppression, and perseverance [15–17]. We explored the effects of inserting a new boundary on model place units. We first familiarized the naive virtual rat with a 65 cm x 65 cm square open environment, during which period the border-grid connectivity and grid-place connectivity self-organized via Hebbian learning. Following this familiarization, the rat explored, without new learning, the familiar environment and a deformed version of this environment containing an additional 40 cm long boundary adjacent to one wall and evenly dividing the space (chosen to match experiment [15]). Again, we observed heterogeneous changes in the time-averaged rate maps of place units (Fig. 3C; grid unit activity depicted in Fig. S1). Some units exhibited place field duplication during boundary insertion, while other units exhibited place field inhibition. Still others persevered largely unaffected. A qualitatively similar mix of responses is observed experimentally during boundary insertions [15–17]. Together, these results demonstrate that many of the heterogeneous place cell behaviors observed across environmental deformations can arise from border cell-grid cell interactions.

### Boundary-tethered grid shifts underlie model grid and place unit distortions

How do model interactions give rise to these grid and place unit distortions? During familiarization, Hebbian learning strengthens the connections from active border units to active grid units at the expense of connections from inactive border units (Fig. 4A; see Materials and Methods). Once familiarized, border unit activity reinstates the grid network state associated with the same pattern of border unit responses during familiarization. This grid reinstatement occurs even when border inputs are activated at a new location, such as when a new or displaced boundary is encountered. In a rescaled open environment, this grid reinstatement leads to ‘shifts’ in the spatial phase of the grid pattern, such that the phase relative to the most recent border input matches the phase entrained during familiarization in the undeformed environment (Fig. 4B,C). Averaged over time (as in Fig. 2A), these *boundary-tethered shifts* can resemble a rescaling of the grid pattern.

**Figure 4.**
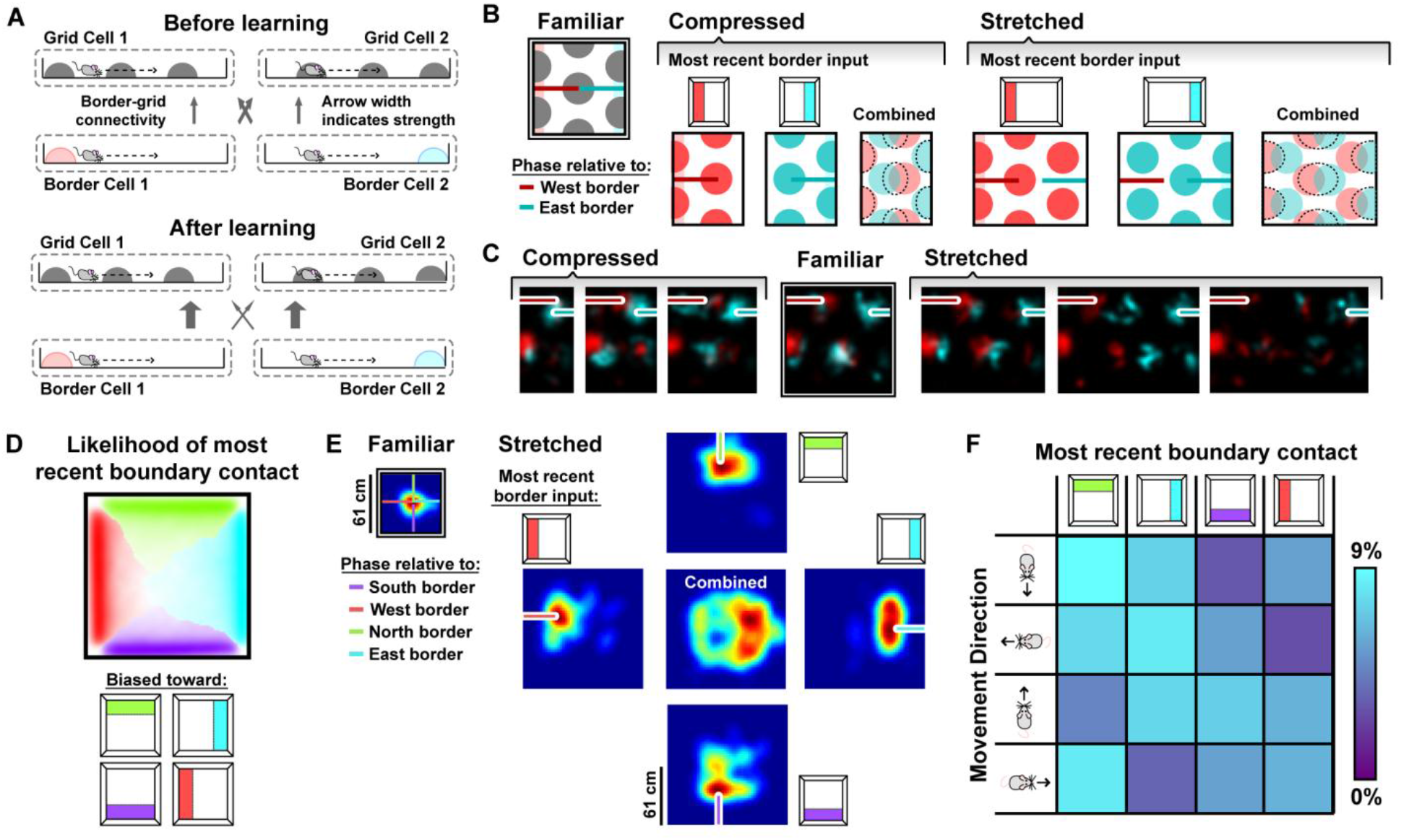
Boundary-tethered grid shifts underlie model grid and place unit distortions. *A) During familiarization, Hebbian learning strengthens the connections between coactive border and grid cells, at the expense of non-coactive connections. B) During deformations, border input acts to maintain the previously learned relationship between grid phase and the most recent border input. C) Rate map of a grid unit following contact with the west border (red), overlaid with the rate map of the same unit following contact with the east border (blue). The spatial phase relative to the most recent border input (indicated by red/blue bars) is preserved during the deformations. Thus the grid pattern is undistorted when separated by the most recent border input. D) Likelihood of having most recently contacted each border as a function of location in a square environment. Hue indicates the most likely recently contacted boundary; saturation denotes the strength of the bias (white – 25% likelihood of sampling; fully saturated – 100% likelihood of sampling). Data from [11]. E) Place fields shift to maintain their familiar relationships relative to the most recent border input. F) Joint probability distribution depicting the relationship between movement direction and the most recently contacted boundary. Data from [11]*.

Why does the appearance of rescaling depend on grid scale and module identity in the boundary-tethered model (Fig. 2A,B)? Because the grid representation is periodic, the border input can only reset the network state to within one period, analogous to a modulo operation. Generally, if the deformation extent is less than the grid period, the different boundaries will reinstate different phases, yielding an apparently rescaled time-averaged pattern. When the deformation extent nearly matches the grid period, different boundaries will reinstate a similar phase, yielding a largely undistorted time-averaged pattern. When the deformation extent exceeds the period, different boundaries will again reinstate different phases; thus the time-averaged pattern will appear distorted. However, in the latter case, additional fields will appear (during stretches) or previously-observed fields will disappear (during compressions). Thus the time-averaged pattern, although distorted, will not resemble a simple rescaling of the grid to match the deformation. Modules are primarily identified by their grid scale -- thus our analysis predicts that the appearance of rescaling will be module-dependent, and that modules with periods less than or equal to the deformation extent will tend not to rescale, consistent with the data in [11]. Furthermore, our model predicts that a grid with a given scale can appear to rescale during less extreme but not during more extreme deformations, consistent with comparison across experiments [9,11,21] (Fig. S2).

Importantly, the likelihood of having most recently encountered a given boundary differs throughout an open environment: locations near a boundary are more likely to be visited following an encounter with that boundary, while central locations are less biased (Fig. 4D). Because of these biases, time-averaged grid fields near a boundary will appear less distorted than central fields during stretching and compression deformations (Fig. 4B,C). Similarly, during partial deformations, locations near the displaced wall are more likely to be visited following contact with it; thus shifts in phase following contact will predominantly affect nearby grid fields, with the phase relationship between this wall and neighboring fields better preserved even after averaging over time (model: Fig. 2C; experiments: [12]). Thus, in this model sampling biases, a product of the particular path of the navigator, mediate the contribution of boundary-tethered shifts to distortions of the time-averaged grid pattern.

A number of theoretical implications follow from the boundary-tethered model. First, this model implies that rescaling and other distortions of the grid pattern are in part an epiphenomenon that results from time-averaging over dynamical shifts in deformed environments. This view offers an alternative to previous accounts that interpret grid rescaling itself as a fundamental phenomenon and propose mechanisms to directly reproduce this effect [19,37]. The boundary-tethered model also implies that environmental deformations induce dynamical shifts in all modules regardless of whether they appear to rescale – this suggests that the appearance or absence of rescaling may not be clear evidence of a functional dissociation between modules [11]. This contrasts with other accounts in which the appearance or absence of rescaling is hypothesized to reflect a fundamental difference in function [11,19].

What about place unit distortions? In this model, place unit activity is constructed as a normalized, thresholded sum of grid unit input [29,30]. Because of the boundary-tethered shifts in grid phase induced during environmental deformations, the location of each place field will also shift, maintaining its spatial relationship to the most recently contacted boundary (Fig. 4E). Critically, as described above, the likelihood of having most recently encountered a given boundary differs throughout an open environment. When averaged across time, these most recent boundary biases result in a mix of place field stretching (closer to displaced boundaries) and bifurcation distortions (further from displaced boundaries). Furthermore, the most recently encountered boundary is correlated with the direction of movement: the rat is more likely to have most recently encountered a given boundary when moving away from it (Fig. 4F). For example, if the rat is traveling eastward in a stretched environment, then the place field will typically be tethered to the west wall; if the rat is traveling westward, then the field will typically be tethered to the east wall. Because the environment has been stretched, west wall-tethered fields will be shifted westward of east wall-tethered fields. Thus, boundary-tethered place field shift causes place fields to be displaced ‘upstream’ along the direction of movement (Fig. 3A). Finally, more extreme deformations of an enclosure lead to more extreme boundary-tethered shifts and less frequent convergence of grid inputs at the same location, and thus systematic decreases in the peak firing rate of place units.

When the rat is trained to run laps on a linear track, movement and likewise the most recently contacted track end are constrained. Thus linear track compressions provide an especially clear view of boundary-tethered updating. Until a track end is encountered, modeled grid and place unit activity unfold according to path integration alone. When a track end is encountered, border input reinstates the grid network state and, in turn, the place network state that coincided with that track end on the familiar track, as seen in Fig. 3B.

Inserting a boundary in an open environment elicits identical border unit activity when either the old boundary or new boundary is nearby in the preferred allocentric direction, inducing boundary-tethered reinstatement of the grid network state at both locations. This grid shift translates to a duplication of the place unit representation adjacent to the old and inserted boundaries. Because a new grid and thus place representation are now active around the inserted boundaries, the old representations previously active at this location in the familiar environment are no longer activated. This leads to an apparent inhibition of place units participating in the old representation (Fig 3C). However, grid and place units that were active at locations distant from the duplicated boundaries will generally persevere unaffected (Fig. 3C).

Thus, in our model, boundary-tethered shifts in grid phase induced by input from border cells drive the diverse grid and place field distortions observed during geometric deformations.

### The predicted boundary-tethered grid shifts are observed in recorded grid cells

Above we have shown that many previously-observed grid and place cell distortions can emerge in part from boundary-tethered shifts in grid phase during environmental deformations. Here, we test whether these shifts can be directly observed in the activity of recorded grid cells during geometric deformations. To this end, we reanalyzed data from two classic environmental deformation studies ([9] and [11]). In [9], rats were familiarized with either a 100 cm x 100 cm square or a 100 cm x 70 cm rectangular open environment, and then reintroduced to deformed and undeformed versions of these environments (i.e. all combinations of chamber lengths and widths of 70 cm or 100 cm), while the activity of grid cells was recorded (familiar square: 23 grid cells; familiar rectangle: 13 grid cells meeting criteria; see Materials and Methods). In [11], rats were familiarized with a 150 cm x 150 cm square open environment, and then reintroduced to deformed (100 cm x 150 cm rectangular) and undeformed versions of this environment, while data were recorded from 51 grid cells.

To test for the predicted boundary-tethered shifts, we first separated the spiking data of each cell according to the most recently contacted boundary, either the north, south, east or west, with contact defined as coming within 12 cm of that boundary [23]. From these data, we created four *boundary rate maps* which summarized the spatial firing pattern of the grid cell after contacting each boundary. Comparison of such rate maps, conditioned on contact with opposing boundaries (north-south vs. east-west), revealed clear examples of grid shift along deformed dimensions (Fig. 5). To quantify shift separately for each dimension, we cross-correlated the opposing boundary rate map pairs (i.e., north-south or east-west boundary pairs). Only pixels sampled after contacting both opposing boundaries were included. Next, we computed the distance from center of the cross-correlogram (0,0 lag) to the peak nearest the center (see Materials and Methods). This distance measures the relative shift between the opposing boundary rate maps. Even in a familiar environment, finite sampling noise will cause this measure of shift to be nonzero. Compared to this baseline, grid shift increased along deformed, but not undeformed, dimensions (combined: Fig. 6A, separated by experiment: Fig. S3A). Moreover, an increase in shift was observed even in cells with small-scale grid patterns which did not rescale (Fig. S4). This indicates that deformation-induced phase shifts affect grid cells even if their time-averaged rate maps do not appear to show rescaling, as predicted by the boundary-tethered model. Note that these shifts were reliably present despite the fact that only approximately one-fourth of the whole-trial data was used to estimate each boundary rate map.

**Figure 5.**
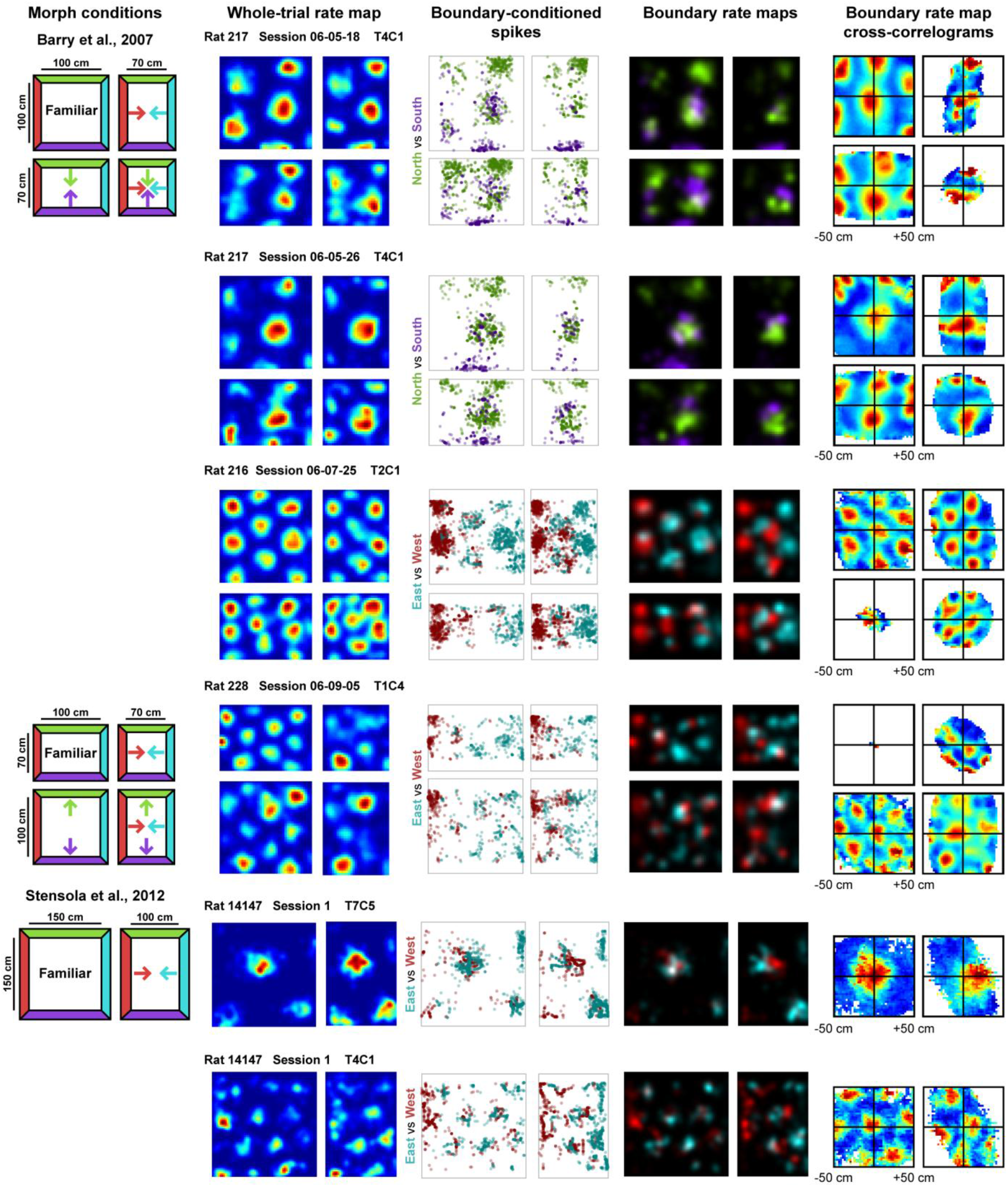
Examples of whole trial rate maps, boundary-conditioned spikes, boundary rate maps, and cross-correlograms of opposing boundary rate maps for recorded grid cells. *Rat, session, and cell identity indicated above whole trial rate maps. Boundary-conditioned spikes and boundary rate maps organized by opposing north-south (green—purple) and east-west (blue—red) boundary pairs. Colored arrows in morph condition indicate the shifts predicted by the boundary-tethered model during each deformation. Note that cross-correlograms only include pixels that were sampled after contacting both opposing boundaries*.

**Figure 6.**
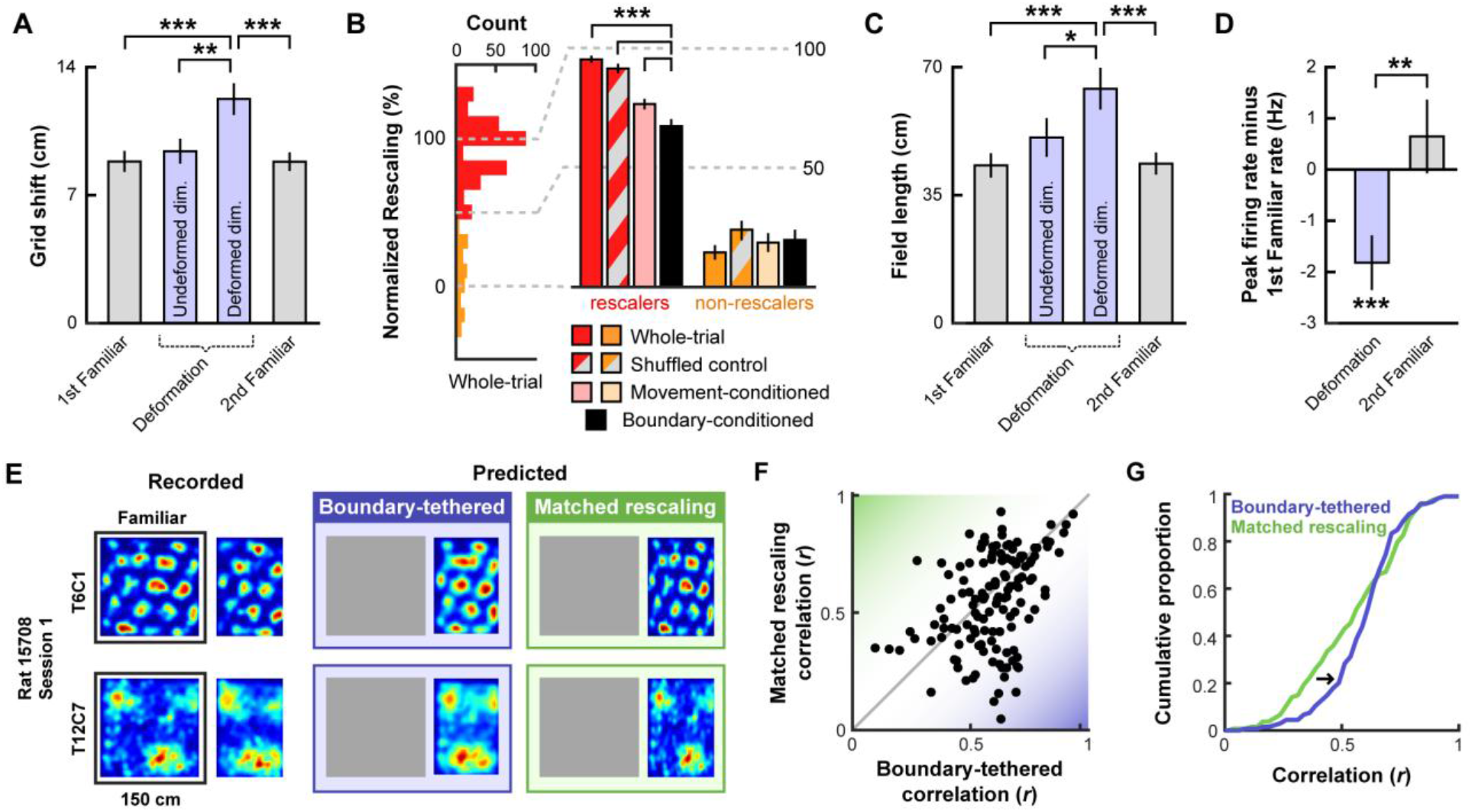
Testing predictions of the boundary-tethered model. All error bars denote mean±SEM. All significance markers denote the outcome of a paired t-test between the indicated conditions. A) Grid shift as measured by the relative phase between opposing boundary rate maps along deformed and undeformed dimensions. (1^st^ familiar vs. deformed: t(80) = 3.98, p < 0.001; undeformed vs. deformed: t(82) = 2.91, p = 0.005; 2^nd^ familiar vs. deformed: t(82) = 4.51, p < 0.001; all other comparisons: t < 1.46, p > 0.148). Data from all experiments in [9,11] combined. B) Whole trial, shuffled control, movement-conditioned and boundary-conditioned grid rescaling factors normalized to range from no rescaling (0%) to a matched rescaling (100%), split by the extent of whole-trial grid rescaling. Because rescaling could vary between simultaneously deformed dimensions within a deformation trial and within cell across deformation trials, rescaling along each deformed dimension and on each deformation trial was included separately (split at 50% rescaling; Boundary-conditioned versus whole-trial, rescalers: t(292) = 11.13, p < 0.001; non-rescalers: t(96) = 1.37, p = 0.173; Boundary-conditioned versus shuffled control, rescalers: t(292) = 8.92, p < 0.001; non-rescalers: t(96) = 0.94, p = 0.349; Boundary-conditioned versus movement-conditioned, rescalers: t(292) = 4.16, p < 0.001; non-rescalers: t(96) = 0.22, p = 0.830). Data from all experiments in [9,11] combined. C) Field length along deformed and undeformed dimensions. (1^st^ familiar vs. deformed: t(80) = 3.70, p < 0.001; undeformed vs. deformed: t(86) = 2.43, p = 0.017; 2^nd^ familiar vs. deformed: t(82) = 3.49, p < 0.001; all other comparisons: t < 1.45, p > 0.151). Data from all compression deformations in [9,11] combined. D) Change in peak firing rate across conditions. (1^st^ familiar vs. deformation: t(80) = 3.57, p < 0.001; 2^nd^ familiar vs. deformation: t(82) = 3.34, p = 0.001; 1^st^ familiar vs. 2^nd^ familiar: t(76) = 0.91, p = 0.364). Data from all experiments in [9,11] combined. E) Examples of recorded and predicted rate maps during one deformation trial for two simultaneously recorded cell from [11]. F) Correlation values between the recorded rate map and the rate maps predicted by the boundary-tethered model versus a matched rescaling. Data from all compression deformations in [9,11] combined. G) Cumulative distribution of the correlation values depicted in (F). The boundary-tethered model results in fewer low-similarity predictions than a matched rescaling indicating a better fit to the experimental data (2-sample Kolmogorov-Smirnov test: D = 0.2030, p = 0.007). *p < 0.05, **p < 0.01, ***p < 0.001.

Next we asked whether the grid pattern in each boundary rate map maintained its spatial phase with the corresponding boundary, as the boundary-tethered model predicts. To address this question, we compared each of the boundary rate maps to the whole-trial familiar environment rate map, while varying the alignment of the two maps along the deformed dimension. If the spatial relationship relative to the most recently contacted boundary is preserved, then each boundary rate map should be most similar to the familiar environment rate map when the two maps are aligned by the corresponding boundary. If, on the other hand, reshaping a familiar environment rescales the grid pattern symmetrically, then the familiar and boundary rate maps should be equally well aligned by either the corresponding or the opposite boundary. Consistent with the boundary-tethered prediction, we found that the correlation between the deformed environment boundary rate map and the familiar environment rate map was maximized when the two maps were aligned by the corresponding boundary rather than the opposite boundary (174 of 246 comparisons; sign test versus 50%: p < 0.001; separated by experiment: Fig. S3B).

The boundary-tethered model further predicts that the appearance of rescaling is in part an epiphenomenon resulting from averaging over trajectories originating from different boundaries. Thus, the appearance of rescaling should be reduced when the data are divided according to the most recently contacted boundary. In contrast, if boundary-tethered shifts did not contribute to the appearance of rescaling, then a similar amount of rescaling should be observed regardless of whether or not data are divided according to the most recently contacted boundary. To test these predictions, we computed the grid rescaling factor between the familiar rate map and each deformed-dimension boundary rate map, aligned by the corresponding boundary. To put this boundary-conditioned rescaling factor into context, we computed three comparison rescaling factors: (1) the classic grid rescaling factor between the familiar rate map and the whole-trial rate map, aligned by the same boundary; (2) a shuffled control in which the grid rescaling factor was computed from a random subset of the whole-trial data, with the amount of data included chosen to match the amount of boundary-conditioned data; (3) a grid rescaling factor conditioned on movement away from the conditioned boundary. This last comparison tests whether changes following boundary-conditioning could alternatively be explained by movement direction, which is correlated with the most recently contacted boundary (Fig. 4F). Boundary-conditioning yielded a significant reduction in normalized grid rescaling factors relative to all three alternative comparisons (combined: Fig. 6B, separated by experiment: Fig. S3C). The reduction in rescaling was specific to cells which previously showed rescaling in their whole-trial rate maps. Thus, boundary rate map grid patterns exhibited significantly less rescaling than whole-trial and movement-conditioned rate maps, consistent with a contribution of border cell-grid cell interactions to the appearance of rescaling.

We next tested whether environmental deformations affect grid field size. The boundary-tethered model predicts that deformations induce shifts in the spatial phase of the grid pattern. Averaged over the entire trial, these shifts should yield an increase in field length primarily along deformed dimensions, regardless of whether the environment was compressed or stretched. On the other hand, a pure rescaling account predicts an increase in field length during stretching, but a decrease in field length during compressions. Because both accounts predict an increase in field length during stretching deformations, we focused on compression trials. From the whole-trial rate maps of each cell we computed the field length during compression deformations, separately along deformed and undeformed dimensions. This analysis revealed an increase in field length along deformed, but not undeformed, dimensions relative to field length in the familiar environment (Fig. 6C), as predicted by the boundary-tethered model. For completeness, we also examined stretching deformations. Field length along deformed dimensions also increased numerically during these deformations (mean ± SEM, familiar: 33.27 ± 5.39 cm; deformed: 34.81 ± 4.17 cm), though this effect did not reach significance in this small sample (n = 13; paired t-test: t(12) = 0.22, p = 0.828).

We then examined firing rate predictions of the boundary-tethered model. If, during deformations, grid vertices are shifted to different locations when different boundaries are encountered, then averaging across trajectories originating from multiple boundaries will necessarily reduce the peak values of the whole trial rate map. Thus the boundary-tethered model predicts a reduction in the *peak firing rate* during environmental deformations, as measured by the peak value of the whole-trial rate map. On the other hand, because the density of grid fields within the environment remains unchanged on average, grid shift does not predict a change in *mean firing rate*, as measured by the total number of spikes across the entire trial divided by the trial duration. Although a pure rescaling account does not make specific predictions about peak and mean firing rates, the simplest assumption would be that neither should change, as the density and intensity of fields tiling the space should be preserved during deformations [38]. Consistent with the predictions of the boundary-tethered model, peak firing rates were significantly reduced during deformation trials relative to familiar trials (Fig. 6D), while mean firing rates did not significantly differ during deformation trials (mean ± SEM, 1^st^ familiar: 2.50 ± 0.24 Hz; deformation: 2.86 ± 0.31 Hz; 2^nd^ familiar: 2.88 ± 0.29 Hz; paired t-test between conditions: 1^st^ familiar vs. deformation: t(80) = 0.54, p = 0.591; 2^nd^ familiar vs. deformation: t(82) = 0.03, p = 0.978; 1^st^ familiar vs. 2^nd^ familiar: t(76) = 0.71, p = 0.479).

Finally, we tested whether deformed rate maps could be accurately predicted by the boundary-tethered model on a trial-by-trial basis. To do so, for each cell and deformation trial we first created predicted boundary rate maps for each displaced boundary from the familiar environment rate map. These rate maps were shifted versions of the familiar rate map, aligned by the corresponding boundary (Fig. S5A). If the length of a boundary changed, then the central portion of the familiar rate map was used to construct the boundary rate map. Next, each boundary rate map was weighted by the actual sampling biases of the rat during that deformation trial. The final boundary-tethered prediction was then the smoothed sum of these weighted predicted boundary rate maps. For comparison, we also computed a rescaled rate map in which the familiar rate map was rescaled to match the deformation. Because additional fields may appear during stretching deformations which were not sampled in the smaller familiar environment, we focused only on compression trials. Across cells, recorded rate maps were more similar to those predicted by the boundary-tethered model than to those predicted by a matched rescaling (Fig. 6E; Fig. S5B), as quantified by the correlations between maps (paired t-test comparing Fisher-transformed correlation values: t(132) = 2.95, p = 0.004; Fig. 6F). This difference was predominately driven by cells whose activity did not resemble a matched rescaling: recorded rate maps which were well-predicted by a matched rescaling were similarly well-predicted by the boundary-tethered model, while recorded maps which were not well-predicted by a matched rescaling were nevertheless well-predicted by the boundary-tethered model. This pattern was reflected in the observation of fewer low-similarity predictions from the boundary-tethered model than from a matched rescaling (Fig. 6G). Thus, the boundary-tethered model can accurately predict individual whole-trial rate maps on a trial-by-trial basis, even when the resulting rate map does not resemble a rescaling.

In sum, we have shown that dividing the grid cell activity according to the most recently contacted boundaries during environmental deformations yields grid patterns which are shifted relative to one another, anchored to the conditioned boundary, and appear less rescaled than the whole-trial grid pattern. Furthermore, we have shown that whole-trial field length increases along deformed dimensions, and whole-trial peak firing rates decrease during deformations while mean firing rate remains unchanged, both matching model predictions. Finally, we have demonstrated that the boundary-tethered model can accurately predict whole-trial rate maps during deformations regardless of whether the resulting maps resemble a matched rescaling. Together, these results provide convergent evidence that boundary-tethered shifts in grid phase contribute to distortions of the grid pattern observed during environmental deformations.

## Discussion

Our results support two primary conclusions. First, many of the complex grid and place cell distortions observed during environmental deformations can emerge from border cell-grid cell interactions. Second, boundary-tethered shifts in grid phase, a hallmark of border cell-grid cell interactions, can be observed directly in the activity of recorded grid cells during deformations. Together, these results highlight previously unrecognized dynamics governing the grid code during environmental deformations and implicate border cell-grid cell interactions as an important contributor to deformation-induced distortions of grid and place cell activity. These results further indicate that time-averaged analyses may have overestimated the malleability of the grid cell spatial metric in response to environmental deformations and suggest that scale-dependent grid rescaling may not be a clear indicator of a functional dissociation between modules. Finally, these results demonstrate that the effects of environmental deformations are not fixed over time, but instead depend crucially on the movement history of the navigator.

A variety of circuits could give rise to boundary-tethered shifts. Here we implemented a particular model of interactions between border, grid and place cells that gave rise to these shifts. This model was feedforward between layers [30], included a path integration-based attractor network of grid cells [28], and generated place cells from grid cell output alone [29]. Although each of these components was motivated by prior work, this model is not intended as a complete recreation of entorhinal-hippocampal connectivity, but rather as a demonstration of how border cell input can give rise to the complex dynamics we describe, even in a relatively simple network. As such, this model excludes known connections that are not essential for these dynamics. For example, this model lacks visual inputs [35], input to place cells from sources other than grid cells [39], and reciprocal connections from place to grid cells [40], all of which play important roles in developing and maintaining a functional spatial code. Moreover, similar boundary-tethered place code dynamics can be observed even before the grid code has fully matured, suggesting that additional mechanisms may contribute to similar dynamics in place cells [41]. Thus, while our results implicate border cell-grid cell interactions as one source of the experimentally-observed grid shifts, additional experiments are required to causally test the particular circuit realization which gives rise to these shifts.

The dynamic boundary-tethered phase anchoring we observe here may reflect a more general phenomenon of grid phase anchoring to external cues or other internal reference frames [8,42]. Consistent with this idea, the grid representation is shaped by a number of boundary and non-boundary cues even in geometrically undeformed environments. For example, grid scale differs between novel and familiar environments [43], the grid pattern is anchored by spatial geometry and other visual features [44,45], and the grid pattern distorts near familiar boundaries as well as in asymmetric environments [44,46]. These effects were not captured by the border cell-grid cell interactions as implemented here, and may reflect phase-anchoring to external cues [8,45,46] or internal reference frames such as boundary vector cells [37,47] or place cells [37,42].

Our results do not rule out additional mechanisms which may be at play during environmental deformations. Indeed, it is likely that multiple mechanisms contribute to the various properties of deformation-induced grid and place field distortions. For example, it is known that during deformations the distorted grid pattern does not persist indefinitely, but relaxes back to the familiar spatial scale with experience [9]. In our simulations, model weights were fixed during deformation trials in order to observe the effects of deformations on model representations free of any obfuscating dynamics. However, even with continued learning, the boundary-tethered model as implemented here cannot capture long-term relaxation dynamics because grid phase and border input are not in conflict long enough for unlearning to occur. More specifically, when the west boundary is encountered following an east boundary contact during an east-west deformation, the border and grid codes are briefly in conflict when the border representation is first activated, causing a small amount of unlearning. However, this border activation also quickly reinstates the learned grid phase, eliminating the conflict between the two. The learned grid phase is then reinforced for as long as the animal remains close to the west boundary, typically long enough to overwrite whatever bit of unlearning had occurred.

Thus, other mechanisms, such as anchoring to additional conflicting reference frames (input from visual cues [8,18,41,48], boundary vector cells [15,22], or place-to-grid feedback [37]) or changes to speed coding [49], are necessary to explain grid relaxation.

Previous work has also revealed conspicuous parallels between deformation-induced distortions of spatial representations in the rat brain and the spatial memory of humans in deformed environments [13,50–52], leading to the suggestion that a common mechanism might underlie these effects. Consistent with this idea, recent evidence suggests that rescaling can be observed in the time-averaged activity of human grid cells [53]. In light of our results, we suggest that boundary-tethered grid shift may be a common mechanism contributing to these cross-species effects, and predict that boundary-anchored shifts in human spatial memory should be observable during environmental deformations.

## Acknowledgements

We are grateful to the laboratories of Edvard and May-Britt Moser, Kate Jeffery and Dori Derdikman for making the data from [11] and [9] available for our reanalysis. We also thank Eli Pollock, Niral Desai and Xuexin Wei for advice on implementing spiking border-grid connections. Finally, we gratefully acknowledge support from nSf grant PHY-1734030 (VB), NIH grants EY022350 and EY022350 (RAE), and NSF IGERT grant 0966142 (ATK). VB thanks the Aspen Center for Physics (Aspen, Colorado; NSF grant PHY-1607611) and the International Center for Theoretical Physics (Trieste, Italy) for support and hospitality while this paper was being completed.

## Materials and Methods

### Model

#### Border layer

The border layer consisted of 32 units. First, the area near each wall in 4 allocentric directions (North, South, East, West) was divided into 8 ‘bricks’ (see [24] for a similar treatment). Each brick extended 12 cm perpendicular from the wall and covered 12.5% of the total environment length along that dimension. Each unit *j* received a uniform input *b_j_* = 0.1 whenever the simulated rat was within one of four adjacent bricks, resulting in a firing field covering 50% of the environment perimeter for each unit. This input was converted to stochastic spiking activity (see below).

#### Grid layer

The grid layer, derived from the model of [28], consisted of 5 grid ‘modules’. Each module consisted of a neural sheet with periodic boundary conditions, visualized as a torus. This neural sheet was composed of 64^2^ identical 2 unit x 2 unit tiles (128^2^ units per module). Each unit in a tile was associated with a particular direction (North, South, East, West), which determined both the movement-direction-specific excitatory input received, as well as its local connectivity. Movement-direction-specific excitatory input *v_j_* to grid unit *j* was determined by

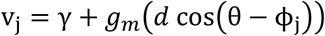

where *d* is the distance moved since the previous timestep, θ is the direction of movement, ϕ_j_ is the preferred direction of unit *j*, *g_m_* is a gain factor specific to the module to which to unit belongs, and γ = 0.6 is a constant. Local connections within each module consisted of shifted radial inhibition, in which each unit inhibited all units within a 12 unit radius by a uniform weight of −0.02. The center of this radial inhibition output for each unit was shifted by 2 units away from that unit in a direction consistent with each units preferred direction. In the absence of other inputs, each grid module yields a hexagonal grid-like pattern of activation on the neural sheet, which is translated during movement at a rate proportional to the gain factor. Thus, to model modules with varying grid scales, the gain factor *g_m_* of module *m* was set by

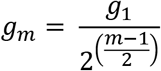

where *g* = 0.45 is the gain of the smallest-scale module, module 1. This results in a geometric series of biologically-plausible [11] grid scales for each module.

#### Place layer

The place layer consisted of 64 units, subject to uniform recurrent inhibition from all place layer units with a weight of −0.15.

#### Border-to-grid connectivity

All grid units received additional excitatory feed-forward projections from all border units. These connections were initialized with random weights uniformly sampled from the range 0 to 0.025, and developed through experience via Hebbian learning (see below and [24]).

#### Grid-to-place connectivity

Each place unit received additional excitatory feed-forward projections from 500 random grid units. These connections were initialized with random weights uniformly sampled from the range 0 to 0.022, and developed through experience via Hebbian learning (see below).

### Model dynamics

#### Activation

The dynamics of the network was developed following the methods in [28]. The activation *a_j_* of unit *j* was determined by first computing the total input *b_j_* to unit *j* according to

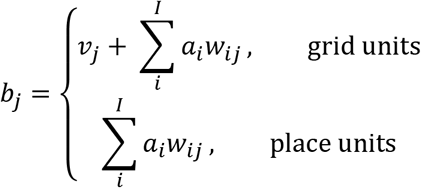

where *a_i_* is a variable quantifying activation of unit *i*, *w_ij_* is the weight from unit *i* to unit *j*, and *I* enumerates all the units. (Note that some weights *w_ij_* can be zero.) Also recall from above that a border unit receives a constant input when the rat is in a boundary region associated with that unit. The total input *b_j_* was used to stochastically determine the spiking *s_j_* of each unit *j* during the current timestep, according to

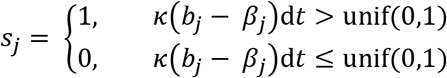

where *κ* = 500 is a scale factor, *β_j_* (border units: *β_j_* = 0; grid units: *β_j_* = 0.1; place units: *β_j_* = 0.05) is the spike threshold for unit *j*, unif(0, 1) is a single draw from a random uniform distribution ranging from 0 to 1, and d*t* = 0.003 sec is the length of each timestep. Finally, this spiking activity was integrated to update the activation variable *a_j_* of unit *j* after each timestep according to

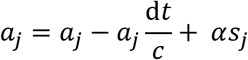

Where *α* = 0.5 is a scale factor and *c* = 0.03 sec is the time constant of integration.

#### Hebbian learning

All Hebbian weights were updated by the competitive learning rule

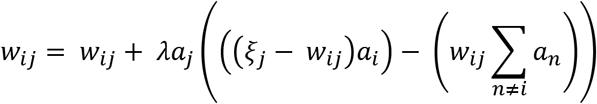

where the sum is only over the set of units with nonzero Hebbian weights to unit *j, λ* = 0.00001 is the learning rate, *ξ_j_* is a constant specific to the connection type (border-to-grid: *ξ* = 0.4; grid-to-place: *ξ* = 0.5) [30,34]. This rule results in competitive activity-dependent weight changes among incoming Hebbian connections, and leads over time to a total weight of *ĵ* across incoming synapses.

### Simulation details

#### Generating simulated rat paths

Because some of the deformed environments that we tested have not been experimentally studied, it was necessary to generate simulated rat paths, rather than using experimentally recorded paths. Open field paths were generated via a bounded random walk model, parameterized by speed and movement direction. At each timestep, unbiased normally-distributed random noise was added to both speed (*σ* = 0.001 cm/msec) and movement direction (*σ* = 1.5 °/msec). To approximate actual rat exploration, speed was bounded to the range [0, 40] cm/sec. If a step would result in the rat path crossing a boundary, random noise was again added repeatedly to the movement direction until the next step would no longer cross the boundary. Open field paths always began in the center of the environment, with the simulated rat stationary and facing a random direction. Linear track paths were generated as straight end to end laps at a constant speed of 20 cm/sec.

#### Familiarization

In all simulations, familiarization with the environment was mimicked by allowing the naive simulated rat to explore the environment for 60 min. Prior to familiarization, grid layer activity was allowed to settle into its grid-like attractor state for 2 sec without learning. Initialization of the grid layer was biased so that an axis of the settled grid network state would lie at an angle of −7.5° relative to east, consistent with experiments [44,46]. Following familiarization, the model weights were saved so that all post-familiarization simulations could begin with the familiarized model.

#### Post-familiarization testing simulations

The model weights were reset to the state saved after familiarization, and the experienced virtual rat was allowed to explore each tested environment for 30 min. Grid layer activity was also initially reset to the familiar environment state corresponding to the rat’s start location. Learning was turned off during the testing phase.

### Analysis

#### Statistical tests

All statistical tests are 2-tailed unless otherwise noted. All error bars denote mean ± 1 standard error of the mean unless otherwise noted.

#### Unit sampling

Due to computational constraints and the redundant nature of grid unit activity, only the spikes from 30 randomly chosen grid units in each module were recorded and analyzed during all simulations. All place units were recorded and analyzed.

#### Rate maps

Rate maps were created by first dividing the environment into 2.5 cm x 2.5 cm pixels. Then the mean firing rate within each pixel was calculated. Finally, this map was smoothed with an isotropic Gaussian kernel with a standard deviation of 1.5 pixels (3.75 cm) and square extent of 9 pixels x 9 pixels (22.5 cm x 22.5 cm). Pixels which were never visited were excluded during further analyses, with the exception of rate map prediction: all pixels were included during rate map prediction as even few missing pixels lead to large gaps of missing pixels following rescaling.

#### Autocorrelations and cross-correlations

*Autocorrelations* of rate maps were computed similar to previous reports [54]. Briefly, the correlation *r* between overlapping pixels of the original rate map and a shifted version of itself was computed as

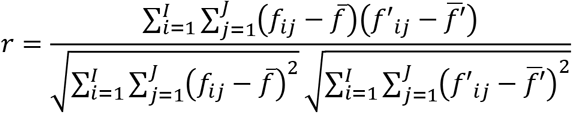

where *f* is the rescaled rate map, *f′* is the familiar rate map, *i* and *j* run over pixels in the overlapping regions of these maps, and 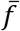 and 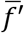 indicate the mean firing rate across overlapping pixels, at a series of single pixel (2.5 cm) step lags. *Cross-correlations* were computed similarly, except that two different rate maps, rather than two copies of the same rate map, were correlated. Autocorrelations and cross-correlations were only estimated for spatial lags with at least 20 overlapping pixels.

#### Grid scale

To compute grid scale for model units we first averaged the autocorrelations of all grid units within a module. Next, we computed the mean distance from the center of the autocorrelation to the center of mass of the six closest surrounding peaks. In cases where the grid period was larger than the size of the environment thus obscuring the periodicity, grid scale was instead estimated by multiplying the scale of the next smaller module by V2, reflecting the parameters set in the attractor model creating the grid. Grid scale for reanalyzed recorded grid cells was computed similarly, but separately from the autocorrelation of each cell.

#### Gridness

To compute gridness for each unit, we first computed the autocorrelation of its rate map and its grid scale. Next we masked the autocorrelation, eliminating all pixels at a distance from the center greater than 1.5 its scale and less than 0.5 its scale. We then computed the correlation between the masked autocorrelation and a rotated version of itself, rotated 30°, 60°, 90°, 120°, and 150°. The final measure of gridness was then the difference between the minimum of the [60° 120°] correlations minus the maximum of the [30° 90° 150°] correlations.

#### Field length

Field length along each dimension was estimated from the autocorrelation by first determining the extent of the central peak of the autocorrelation, defined as all contiguous pixels with correlation values greater than 10% of the maximum correlation. Next, field length was computed separately for each dimension as the distance between the most extreme pixels within this central peak along that dimension.

#### Grid rescaling facto

The grid rescaling factor during each deformation trial was computed separately for each unit by comparing rescaled versions of the familiar environment rate map to the deformed environment rate map. Following [11], the familiar rate map was uniformly rescaled to a series of chamber lengths, ranging from 10 cm below the smaller of the deformed and familiar chamber lengths, through 10 cm above the larger of these chamber lengths in 5 cm (2 pixel) increments. This yielded a set of rescaled familiar rate maps for each unit. For each rescaled map, we computed the correlation r (defined above) between the deformed and rescaled rate maps twice, once when the two rate maps were aligned by each opposing boundary. The grid rescaling factor was then defined as the ratio between the rescaled chamber length that yielded the highest correlation and the familiar chamber length, across either alignment. When comparing rescaling factors between whole-trial and boundary-conditioned data, rescaling was only computed for alignment by the conditioned boundary.

#### Grid shift analysis

To test these data for the presence of grid shifts during environmental deformations, we first divided the spiking activity of each cell according to the most recent boundary contact (North, South, East, or West). Boundary contact was defined as the rat being within 12 cm of a boundary. Spiking activity prior to boundary contact at the beginning of the trial was ignored. Next, four separate rate maps were created, one for each most recently contacted boundary. To quantify grid shift along a particular dimension for each cell, the rate maps of opposing boundaries perpendicular to the chosen dimension were cross-correlated at a series of lags in single pixel steps (see above) within the range of ±20 pixels (±50 cm). Only pixels sampled after contacting both opposing boundaries were included in these cross-correlations. The distance from the center to the nearest peak of this cross-correlogram was computed as the measure of grid shift. The nearest peak was defined by first partitioning the cross-correlogram into ‘blobs’ of contiguous pixels which had correlations of at least 30% of the maximum value. Then, the location with the maximum correlation value within the blob nearest to the center was taken as the nearest peak.

#### Reanalysis of experimental data

A complete description of the experiments was provided in [9,11]. Data from [9] included an initial set of 66 putative cells, from which 38 cells meeting various criteria were selected as grid cells for analysis in the original publication. Similarly, we included only cells with average gridness across both familiar trials >0.4 from this dataset, yielding 36 included grid cells. Note that unlike in [9] we did not exclude cells which were poorly fit by rescaling during deformation trials, as the boundary-tethered model predicts that distortions which do not resemble a rescaling may occur. For alignment, rescaling, and rate map prediction analyses, first familiar trial rate maps were used for comparison; in the few cases where no rate map was recorded during the first familiar trial, the rate map from the second familiar trial was used instead.

#### Boundary-tethered rate map prediction

For each cell and deformation trial we first created predicted boundary rate maps for displaced boundaries from the familiar environment rate map. These rate maps were shifted versions of the familiar rate map, aligned by the corresponding boundary (Fig. S5A). If the length of a boundary changed, then the central portion of the familiar rate map was used to generate the predicted boundary rate map. Next, sampling biases were applied as follows. First, a map of the actual sampling behavior following each boundary contact during the deformation trial was computed, as described in the ‘rate maps’ section above. From these maps the probability of having most recently contacted each boundary was computed at each pixel. The contribution from each boundary rate map was then weighted by this probability. The final rate map predicted by the boundary-tethered model was then the sum of these weighted boundary rate maps, smoothed with the Gaussian kernel described in the ‘Rate maps’ section above.

#### Data and code availability

All simulations were conducted with custom-written MATLAB scripts. These scripts and the simulation results presented here are available from the authors upon request. All reanalyzed data are available upon request from the corresponding authors of the relevant papers.

---

**Supplementary Figure 1.**
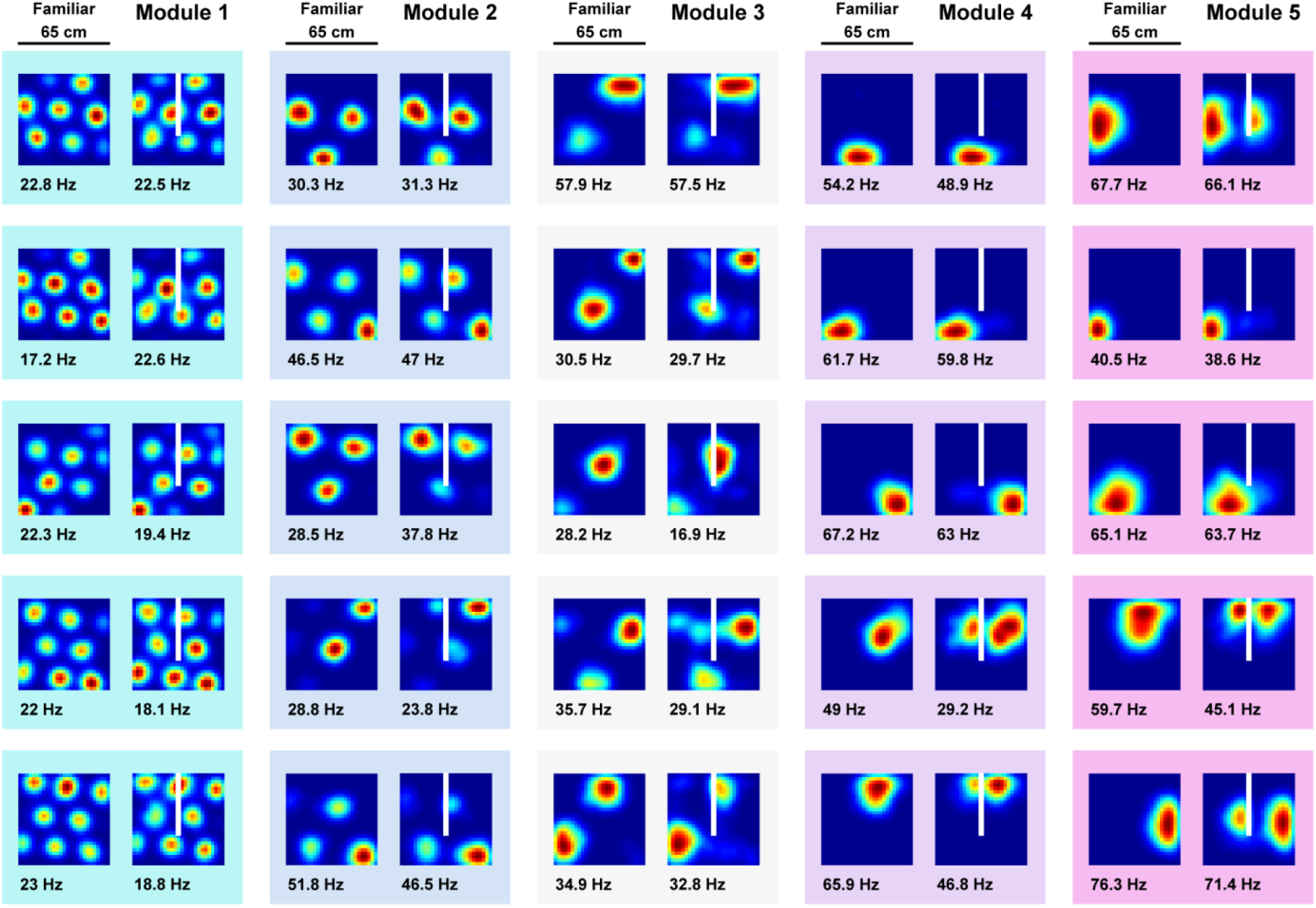
Grid unit activity during insertion of a new boundary in an open environment. Examples of whole-trial grid unit activity during exploration of a familiar chamber and boundary insertion (white line) – five random units shown from each module. Distortions are minimal in the time-averaged rate maps of small-scale grid units (as observed experimentally [20]), but become apparent in the activity of large-scale grid units. Peak firing rate noted below the lower left corner of each map. Color normalized to the maximum for each rate map.

**Supplementary Figure 2.**
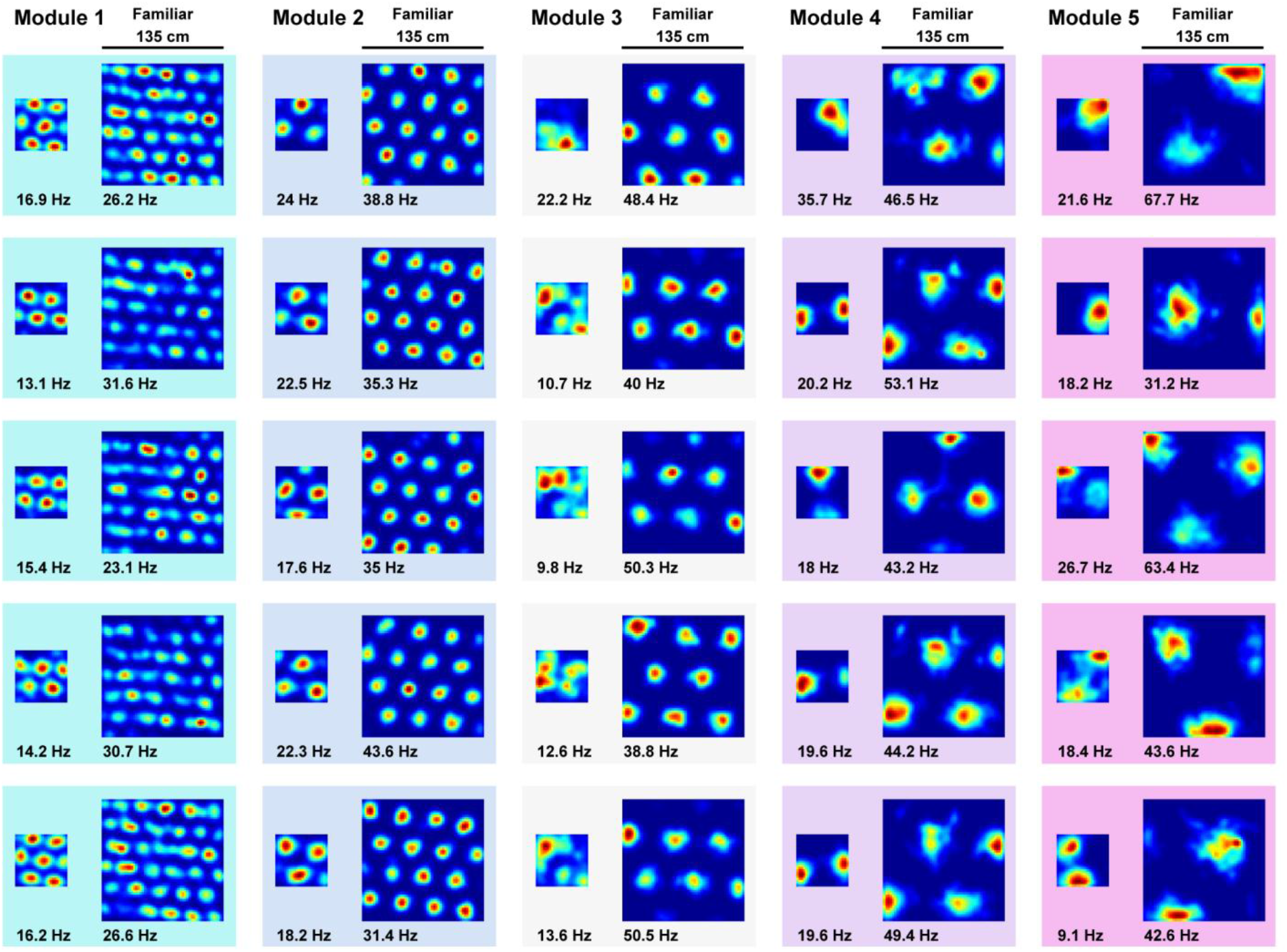
Model grid units do not rescale during a more extreme compression deformation. Although grid rescaling was reported during deformation in two electrophysiological studies [9,11], another study implementing a more extreme compression deformation experiment did not report evidence of rescaling in grid cells [21]. To test whether the boundary-tethered model could account for a lack of rescaling during this more extreme compression, we familiarized the naïve virtual rat with a 135 cm x 135 cm square environment. After this familiarization, the rat then again explored the familiar environment and a compressed 58 cm x 58 cm version of this environment without new learning. During this extreme compression, model grid units did not resemble a rescaling, replicating experimental observation. Five random grid units from each module, peak firing rate denoted in bold below each map. Color normalized to the maximum for each rate map.

**Supplementary Figure 3.**
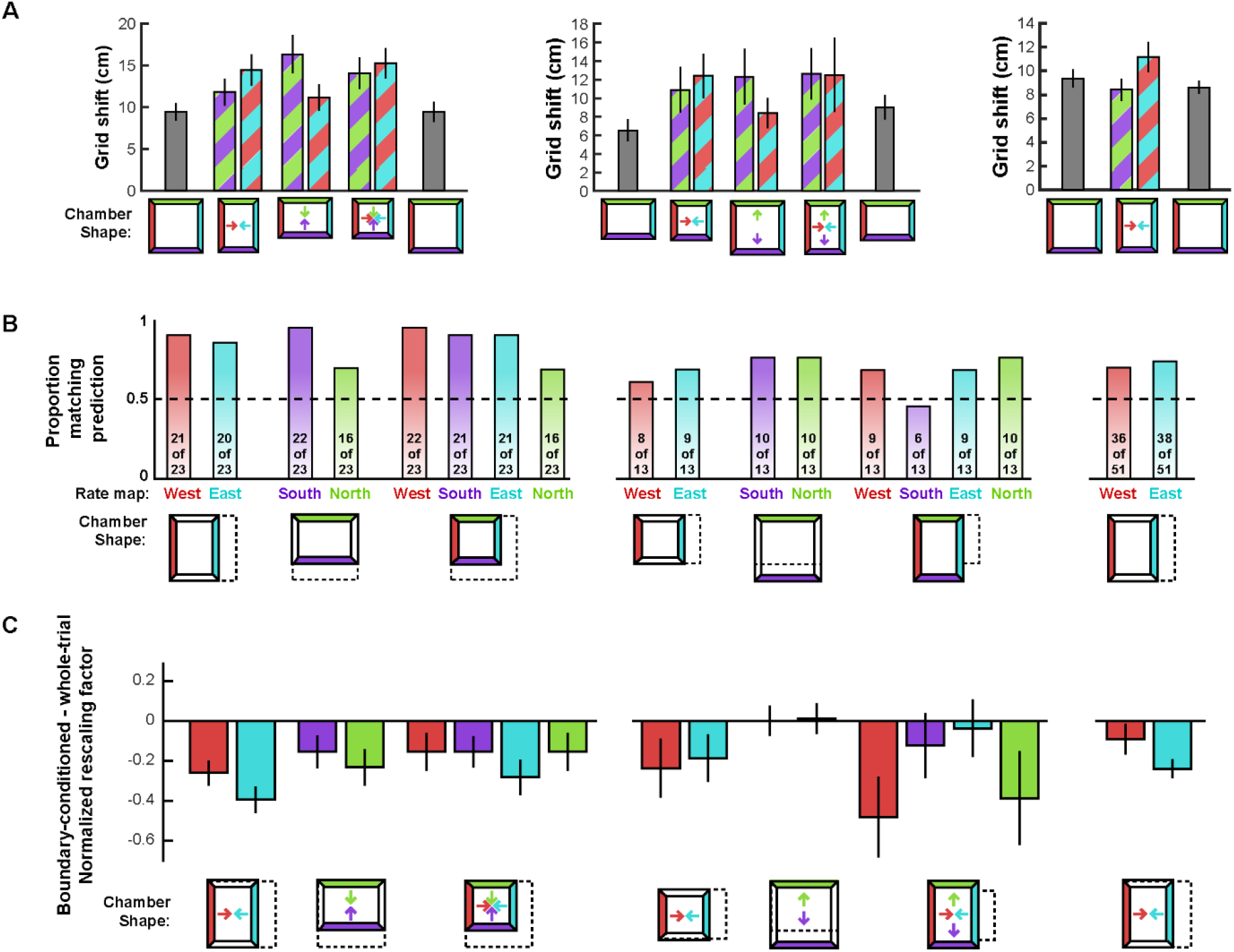
Grid shift, alignment, and boundary-conditioned rescaling of recorded grid cells separated by condition. In each case rats trained in (left) a familiar square (data from [9]), (middle) a familiar rectangle (data from [9]), and (right) a familiar square (data from [11]). A) Grid shift computed for each condition separately (see Text; errors bars ± 1 SEM). Colored arrows indicate the dimensions along which our model predicts an increase in shift above baseline grid shift. B) Proportion of trials for which each boundary rate map was best matched with its familiar environment rate map when aligned by the most recently contacted boundary (as predicted by the boundary-tethered model) versus the opposing boundary (counts shown within the bars). Familiar environment (dashed box), deformed environment (solid walls), and boundary (colored walls) shown in lower insets (familiar and deformed environments aligned by arbitrary walls to make the change in shape apparent). C) Change in normalized rescaling factors following boundary-conditioning separately for each condition (boundary-conditioned minus whole trial; errors bars ± 1 SEM).

**Supplementary Figure 4.**
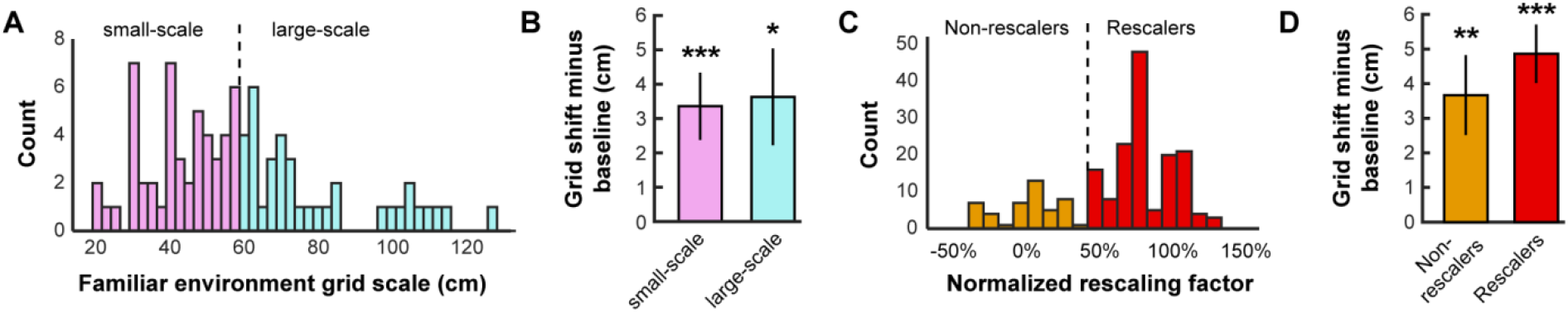
Grid shift is observed in small-scale and non-rescaling recorded grid cells. Data from all experiments [11] and [9] combined. A) Histogram of grid scales averaged across familiar trials. B) Grid shift along deformed dimensions after subtracting average shift during familiar trials. A significant increase in grid shift above familiar baseline was observed for small-scale (grid scale < 60 cm; paired t-test versus familiar shift: t(51) = 3.55, p < 0.001) and large-scale grid cells alike (t(34) = 2.64, p = 0.012), with no significant difference between conditions (2-sample t-test: t(85) = 0.17, p = 0.866). C) Histogram of normalized grid rescaling factors. Grid rescaling normalized such that no rescaling corresponds to 0% and rescaling completely to match the deformation corresponds to 100%. Because rescaling could vary between simultaneously deformed dimensions within a deformation trial and within cell across deformation trials, rescaling along each deformed dimension and each trial was included separately. D) Grid shift along deformed dimensions after subtracting average shift during familiar trials. As in (C), grid shift along each deformed dimension and each trial was included separately. A significant increase in grid shift above familiar baseline was observed in rescalers (normalized rescaling factor ≥ 50%; paired t-test versus familiar shift: t(131) = 6.02, p < 0.001) and non-rescalers (t(61) = 3.274, p = 0.002) alike, with no significant difference between conditions (2-sample t-test: t(192) = 0.85, p = 0.397).

**Supplementary Figure 5.**
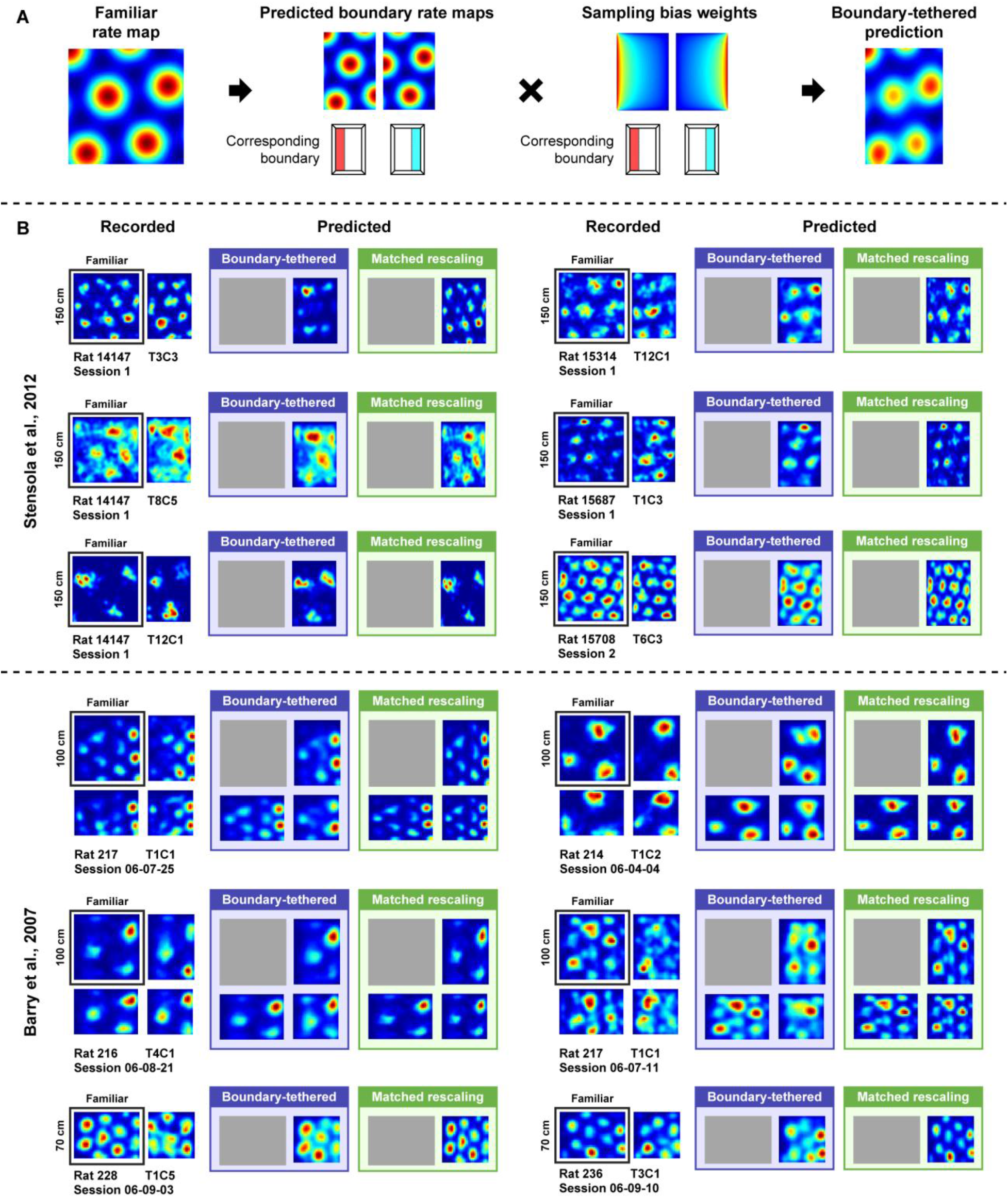
Predicting whole-trial rate maps with the boundary-tethered model. A) To predict rate maps from the boundary-tethered model for each cell and compression deformation trial we first created predicted boundary rate maps from the familiar environment rate map for each displaced boundary. These rate maps were shifted versions of the familiar rate map, aligned by the corresponding boundary. If the length of a boundary changed, then the central portion of the familiar rate map was used to construct the boundary rate map. Next, the contribution of each boundary rate map at each location was weighted by the actual probability of sampling that location following contact with the corresponding boundary for that deformation trial, computed from the actual path of the rat during that deformation trial. The final boundary-tethered prediction was then the smoothed sum of these predicted boundary rate maps. B) Example recorded rate maps, accompanied by the predictions from the boundary-tethered model and a rescaling matched to the extent of the deformation. Rat, session, and cell identity indicated below each set of recorded rate maps.

## Notes

Conflict of interest: The authors declare no competing conflicts of interest.

